# Alcohol Dependence Differentially Alters Action and Outcome Related Orbitofrontal Cortex Activity

**DOI:** 10.1101/2020.07.03.186148

**Authors:** Christian Cazares, Drew C. Schreiner, Christina M. Gremel

## Abstract

Alcohol dependence results in long-lasting deficits in decision-making and behavioral control. Neurobiological investigations have identified orbitofrontal cortex (OFC) as important for value contributions to decision-making as well as action control, and alcohol dependence induces long-lasting changes to OFC function that persist into protracted withdrawal. However, it is unclear which contributing OFC computations are disrupted in alcohol dependence. Here, we combined a well-validated mouse model of alcohol dependence with in vivo extracellular recordings during an instrumental task in which lever press duration serves as the contingency, and lever pressing is sensitive to outcome devaluation. We found prior alcohol dependence did not impair use of duration contingency control but did reduce sensitivity to outcome devaluation. Further, alcohol dependence increased OFC activity associated with lever-pressing but decreased OFC activity during outcome-related epochs. Hence, alcohol dependence induces a long-lasting disruption to OFC function such that activity associated with actions is enhanced, but OFC activity in relation to outcomes is diminished. This has important implications for hypotheses regarding compulsive and habitual phenotypes observed in addiction.

## Introduction

Alcohol Use Disorder (AUD) can result in decision-making deficits that can persist into protracted abstinence. In those suffering from AUDs, these deficits are thought to contribute to habitual and compulsive alcohol-seeking, a persistent vulnerability to relapse, and decrements in daily cognitive function (Stephens & Duka, 2008; Berre et al., 2012; Reich & Goldman, 2015; Le Berre et al., 2017; Sebold et al., 2017; Bickel et al., 2018). With reports of alcohol dependence-induced functional and structural alterations across the cortex (Volkow et al., 1994, 1997; Laakso et al., 2002; Cardenas et al., 2011; Durazzo et al., 2011; Beck et al., 2012; Sjoerds et al., 2013; Thayer et al., 2016), it is highly likely that a broad array of computations normally contributing to efficacious decision-making are also altered. Identifying which computations are disrupted along with any corresponding aberrant activity patterns would offer a starting point for mechanistic investigations into cortical circuit alterations that produce these long-lasting impairments in decision-making.

One such cortical circuit that often shows long-lasting dependence-induced disruptions is the orbital frontal cortex (OFC). Abstinent AUD patient studies generally report a hypoactive OFC both at baseline and during adaptive decision-making (Volkow et al., 1994, 1997; Boettiger et al., 2007; Sjoerds et al., 2013; Reiter et al., 2016), but also report OFC hyperactivity to stimuli and related approach behaviors (Wrase et al., 2002; Hermann et al., 2006; Reinhard et al., 2015), reminiscent of OFC hyperactivity reported in patients with other psychiatric conditions, including obsessive compulsive disorder (Milad & Rauch, 2012; Pauls et al., 2014; Robbins et al., 2019; Lüscher et al., 2020). This dichotomy of effects suggests that long-lasting perturbations to OFC circuitry induced by alcohol dependence may differentially alter the computations performed by OFC neurons in response to information coming into OFC. Hence, initial investigations into computations performed by OFC during decision-making and their long-lasting disruption in alcohol dependence would provide a framework with which to investigate broader circuit mechanisms contributing to observed OFC dysfunction.

Several lines of evidence implicate the OFC as a key contributor to computations that can contribute to value-based decision-making processes (Fellows, 2007; Wallis, 2007; Gremel & Costa, 2013; Stalnaker et al., 2015; Padoa-Schioppa & Conen, 2017) as well as to compulsive control (Milad & Rauch, 2012; Pauls et al., 2014; Robbins et al., 2019; Lüscher et al., 2020). In particular, manipulations of OFC function support a role for OFC in the updating and retrieval of expected outcome value to control behavior (Gremel & Costa, 2013; Rhodes & Murray, 2013; Baltz et al., 2018; Malvaez et al., 2019), as well as in compulsive action control (Ahmari et al., 2013; Burguière et al., 2013; Pascoli et al., 2015, 2018). Animal models of alcohol dependence have revealed long-lasting dependence-induced disruptions to OFC dependent processes including behavioral flexibility and outcome devaluation (Badanich et al., 2011; Kroener et al., 2012; M. F. Lopez et al., 2014; Fernandez et al., 2017; Renteria et al., 2018, 2020). Alcohol dependence results in long-lasting changes to OFC intrinsic excitability and synaptic transmission, as well as increases in dendritic spine density of OFC neurons (Badanich et al., 2013; McGuier et al., 2015; Nimitvilai et al., 2016; Nimitvilai, Lopez, et al., 2017; Nimitvilai, Uys, et al., 2017; Renteria et al., 2018). We previously reported that alcohol dependence led to changes in OFC excitability as well as an insensitivity to outcome devaluation, as reflected by habitual control, in protracted withdrawal. Notably, artificially increasing the activity of OFC projection neurons was sufficient to restore sensitivity to outcome devaluation in alcohol dependent mice (Renteria et al., 2018). To this end, the observed dependence-induced deficits in decision-making are hypothesized to include a breakdown of critical OFC computations.

Here, we examined dependence-induced long-lasting disruptions to OFC computations performed during protracted withdrawal using an instrumental task in which actions were made for food outcomes. To do so, we adapted an action contingency task, historically termed action differentiation (Platt et al., 1973; Kuch, 1974; Yin, 2009; Fan et al., 2012). In this task, mice must learn to press and hold a lever down beyond a fixed minimum duration to earn a food reward; hence, duration serves as the contingency. Outcome devaluation testing procedures followed initial task acquisition. Prior works have suggested that alcohol dependent rats and mice show largely similar acquisition of lever-press performance compared to naïve controls, but outcome devaluation procedures (Corbit et al., 2012; M. F. Lopez et al., 2014; Morisot et al., 2019; Renteria et al., 2018, 2020), and more recently, contingency degradation procedures (Barker et al., 2020), have shown lever pressing is under habitual control. We found that daily performance in our contingency task was largely intact in alcohol dependent mice; however, subsequent devaluation testing recapitulated the dependence-induced reduced sensitivity to outcome devaluation. Examining OFC activity during acquisition of this task, we found that prior induction of alcohol dependence led to higher OFC firing rates related to lever-pressing, but reduced firing rates during periods associated with outcome retrieval.

## Results

### Induction of ethanol dependence disrupts goal-directed control over decision-making

We employed a well-validated model of chronic intermittent ethanol (CIE) vapor exposure and repeated withdrawal (H. C. Becker & Hale, 1993; Howard C. Becker & Lopez, 2004; Marcelo F. Lopez & Becker, 2005; Griffin et al., 2009). Mice were exposed to periods of ethanol (CIE) or air (Air) vapor and subsequent withdrawal over a period of four weeks (one vapor cohort; Air: n = 8, CIE: n =7) (Figure 1A). CIE procedures produced mean blood ethanol concentrations in ethanol-exposed mice in line with previous reports (29.53 ± 2.36 mM) (Marcelo F. Lopez & Becker, 2005; Renteria et al., 2018). Alcohol withdrawal has been delineated into two phases; an immediate acute withdrawal period (2-3 days), followed by a protracted period extending at least three months (Heilig et al., 2010). To examine OFC activity and related behavior during this protracted withdrawal period, food-restricted mice began instrumental training and testing procedures 5 days after their last vapor exposure.

**Figure 1.**
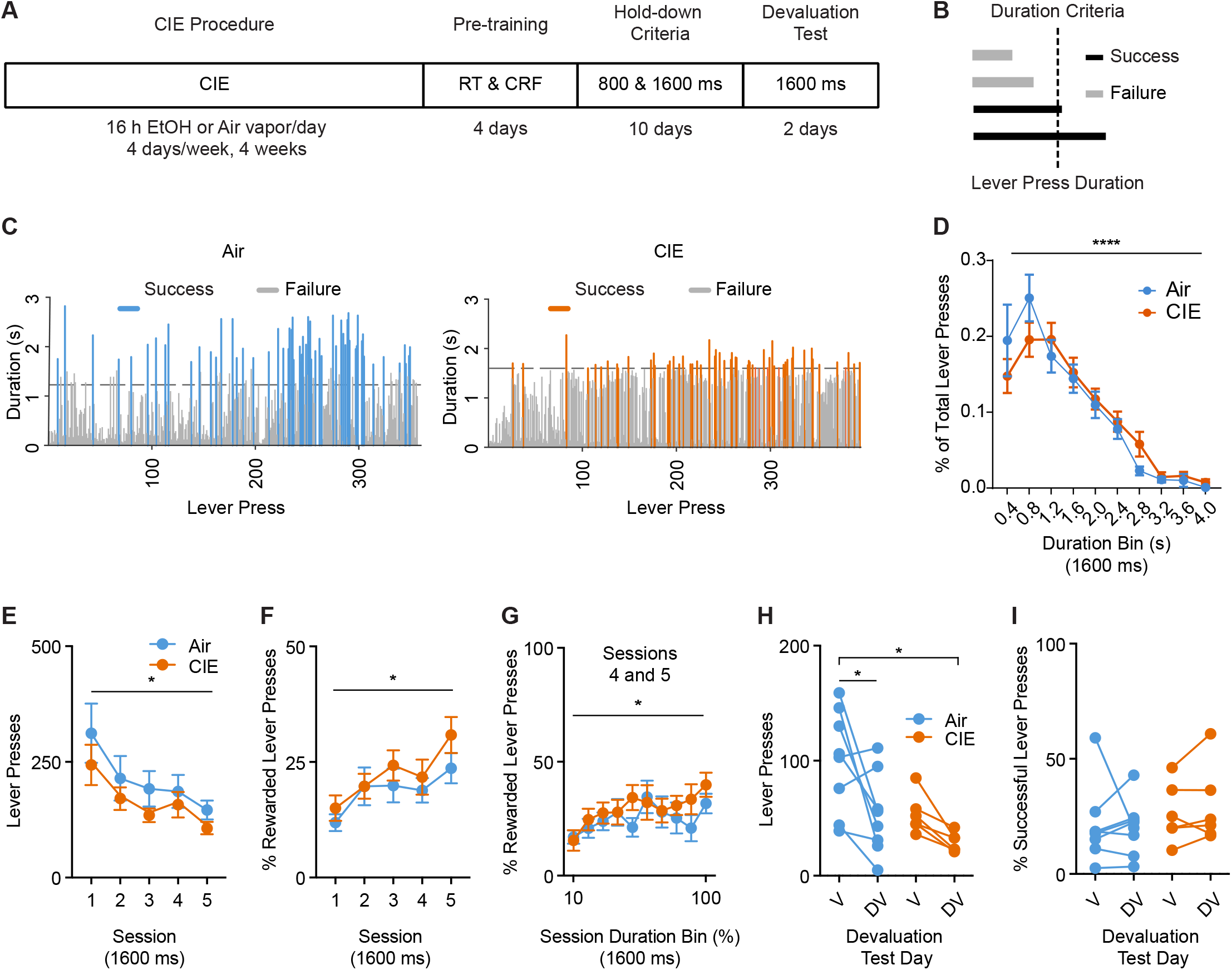
Effects of alcohol dependence on action differentiation and outcome devaluation. **(A).** Experimental timeline starting with the CIE procedure and subsequent Random Time (RT) and Fixed-Ratio Continuous Reinforcement (CRF) schedules of reinforcement, followed by 5 daily sessions of 800-ms lever press duration criterion sessions and 5 daily sessions of 1600-ms lever press duration criterion sessions, and 2 subsequent days of outcome devaluation (DV) testing. **(B).** Schematic of lever press duration performance. Lever presses exceeding the session’s minimum hold-down duration criterion were rewarded only at the offset of the lever press. **(C).** Example lever press performance from individual Air and CIE mice occurring within a single 1600-ms hold-down duration criterion session late in training. **(D).** Distribution of lever press durations, **(E).** average total lever presses, **(F)** and average percentage of rewarded lever presses through the 5 daily 1600-ms lever press duration criterion sessions. **(G).** Average percentage of rewarded lever presses across normalized session duration bins during the last two days of 1600-ms hold down criterion training sessions. **(H).** Average total lever presses in Valued and Devalued states throughout DV testing. **(I).** Average percentage of successful lever presses in Valued and Devalued states throughout DV testing. Data points represent mean ± SEM. *p<0.05, ****p<0.0001.

To examine the effects of prior CIE procedures on OFC activity during action and outcome-related epochs in decision-making, we adapted an instrumental task examining action differentiation (Figure 1A), where a mouse must learn to press and hold a lever down beyond a fixed minimum duration to earn a reward (Yin, 2009; Fan et al., 2012). Throughout training, mice learned to press and hold down a lever beyond a pre-determined minimum duration in order to earn a food pellet upon release of the lever. Mice self-initiated and self-terminated every lever press in the absence of any extrinsic cues signaling lever press duration. Importantly, reward delivery occurred only at the offset of a lever press that exceeded the duration criteria (Figure 1B and 1C), preventing the use of reward presence to signal lever press termination. Further, this task produces discretized behavior epochs conducive to neural activity analysis (e.g. lever-press onset, during the lever-press, lever-press offset, and outcome delivery).

Following vapor procedures, and after initial lever press training (see Materials and Methods), Air and CIE mice underwent action differentiation training, with the initial criterion for lever press duration set at 800-ms for 5 daily sessions, followed by 5 daily sessions of a 1600-ms criterion (see Materials and Methods). Representative sessions from one Air and one CIE mouse on a 1600-ms criteria day are shown in Figure 1C, suggesting that mice show a distribution of lever press durations. This distribution of lever presses was similar between Air and CIE mice (1600-ms criteria training, two-way repeated measures ANOVA (Bin x Treatment); no interaction; main effect of Bin: F(9, 117) = 30.88, p<0.0001), and each group showed a rightward shift in the distribution following a switch from 800-ms to 1600-ms training criteria (Figure 1 — figure supplement 1A and 1B). We examined behavior throughout the 1600-ms criteria sessions, after mice had learned the action differentiation rule and had shifted to a longer duration contingency. Air and CIE mice showed similar levels of lever pressing that increased across sessions (two-way repeated measures ANOVA (Session x Treatment); no interaction; main effect of Session only: F(4, 52) = 9.77, p<0.0001) (Figure 1E). As reflected in the duration distribution of the representative sessions (Figure 1C), only ~25% of lever presses in Air and CIE mice exceeded the duration criterion within a session (two-way repeated measures ANOVA (Session x Treatment); no interaction; main effect of Session: F(4, 52) = 8.72, p<0.0001) (Figure 1F). While we did see a slight difference in response rates between groups (Figure 1 — figure supplement 1C), when we examined whether CIE exposure produced different reward rates within a session, we found similar levels of rewarded lever presses between groups (Figure 1G). To investigate the efficiency of actions throughout a session, we compared the average proportion of total lever presses in well-trained Air and CIE mice that exceeded the 1600-ms duration criterion within equidistant duration bins throughout the last two sessions. A two-way repeated measures ANOVA (Bin x Treatment) found no interaction, and a main effect of Bin only: F(9, 144) = 3.11, p=0.002, suggesting a similar reward rate across a session. Thus, CIE mice and Air controls acquire similar levels of lever pressing for food, with CIE mice showing no impairments in using duration as the contingency to gain the outcome.

To examine whether lever pressing was controlled by the expected outcome value in Air vs. CIE mice, we used an outcome devaluation procedure following training on the 1600-ms duration criterion. In outcome devaluation testing, a reduction in lever-pressing in the devalued state compared to valued state reflects decision-making sensitive to the expected outcome value (Dickinson, 1985). Prior work has found that alcohol dependence reduces the contribution of expected value to decision-making (Dickinson, 1985; M. F. Lopez et al., 2014; Renteria et al., 2018), and we performed this testing within a time period (21 days following the last vapor exposure) when reduced value control has previously been observed (Renteria et al., 2018). We subjected Air and CIE mice to sensory-specific satiation of food pellets previously earned by lever pressing or to a previously habituated control outcome (20% sucrose solution). In each of the two consecutive test days, mice had ad-libitum access to either pellets (devalued state) or sucrose solution (valued state) for 1 hour after which we measured non-reinforced lever press responses. We performed a two-way repeated measures ANOVA (Value x Treatment) to examine whether treatment groups showed different patterns of responding. We found main effects of Value (F(1, 12) = 10.45, p=0.01) and Treatment (F(1, 12) = 7.133, p=0.02) (no interaction), suggesting a difference in the level of lever pressing between groups during testing and greater overall responding on valued than devalued days. To test our *a priori* hypothesis that CIE procedures would reduce the sensitivity to outcome devaluation procedures (i.e. a reduced magnitude effect), pre-planned post-hoc comparisons showed a significant difference between devalued and valued lever pressing only in Air animals (p=0.01), but not CIE mice (p=0.31) (Figure 1H). In contrast, the proportion of successful lever presses that exceeded the duration criteria remained similar across devaluation states for both treatment groups (p>0.3) (Figure 1I). As Air and CIE mice consumed similar amounts during the pre-feeding period on the devalued day (Figure 1 — figure supplement 1D), the present data provide additional evidence that prior CIE exposure induces a long-lasting reduced sensitivity to outcome devaluation and suggests that computations normally supporting such decision-making may be altered.

### OFC populations differentially encode decision-making task components

Given that the deficits in outcome devaluation observed in CIE mice have been shown to involve the OFC (Gourley et al., 2013; Gremel & Costa, 2013; Rhodes & Murray, 2013; Gremel et al., 2016; Renteria et al., 2018), and OFC modulates activity during outcome-related epochs (Rolls et al., 1996; Wallis, 2012), we hypothesized that alcohol dependence would disrupt OFC neural activity related to outcome-related epochs in our task. We implanted additional cohorts of mice with chronic indwelling micro-electrode arrays into the OFC prior to the start of vapor procedures and action differentiation lever press training (five vapor cohorts; Air n = 9, CIE n = 9) (Figure 2A). The similarity of daily performance between Air and CIE mice, as observed in the behavioral cohort, allowed us to compare associated neural activity in the absence of any gross behavioral performance differences (Figure 2 — figure supplement 1).

**Figure 2.**
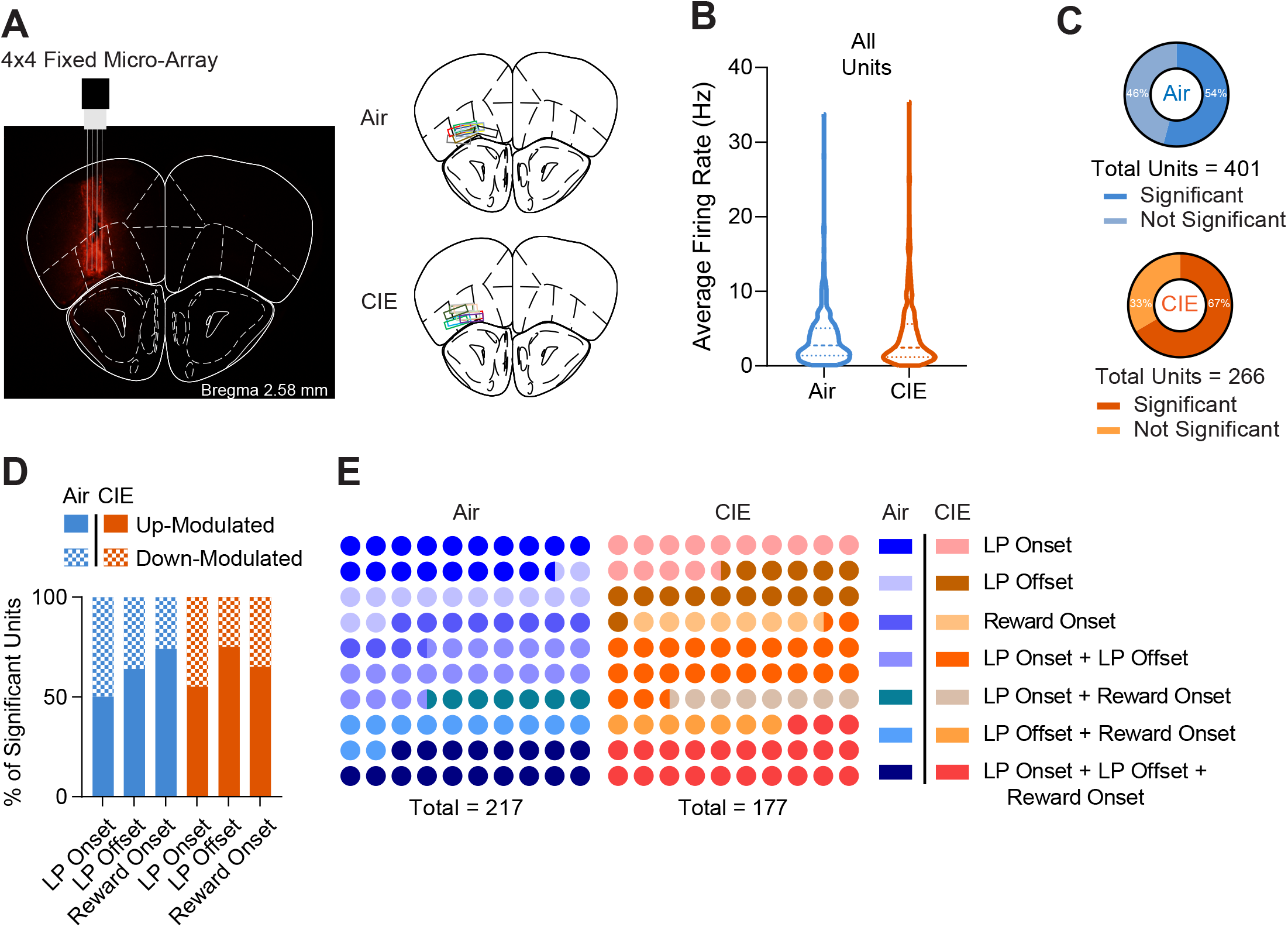
OFC activity correlates of actions and outcomes. **(A).** Representative image of fixed micro-array implant over the orbitofrontal cortex. Implant locations segmented by Air and CIE groups. A subset of micro-arrays was dyed with a 25 mg/ml Dil (1,1’-Dioctadecyl-3,3,3’,3’-tetramethylindocarbocyanine perchlorate) solution in 200 proof ethanol for placement verification. **(B).** Average firing rates from all captured units during a baseline period (−5 to −2 seconds before lever press onset) for Air and CIE cohorts. **(C).** Proportion of all captured units in which firing rates significantly deviated from a baseline period (−5 to −2 seconds before lever press onset) for Air (~54%) and CIE (~66%) groups. **(D).** Percentage of units that significantly increased or decreased their firing rates from baseline in relation to lever press onset, offset, and food pellet reward delivery. **(E).** Proportion of units that significantly changed their firing rates from baseline in relation to discrete task events (Lever Press Onset: Air: ~18%, CIE: ~15%; Lever Press Offset: Air: ~13%, CIE: ~16%; Reward Delivery: Air: ~11%, CIE: ~7%) as well as multiple task components (Lever Press Onset and Offset: Air: ~20%, CIE: ~24%; Lever Press Onset and Reward Delivery: Air: ~6%, CIE: ~7%; Lever Press Offset and Reward Delivery: Air: ~12%, CIE: ~7%; Lever Press Onset, Offset, and Reward Delivery: ~18%, CIE: ~23%).

We focused on OFC activity data collected during the last two sessions of the 1600-ms duration criterion, a time-point during which animals from both groups most proficiently performed the task. Putative single OFC unit spike activity was aligned to timestamps collected each time a lever press onset, offset, or pellet reward delivery occurred (see Materials and Methods). Importantly, there was no effect of CIE exposure on average baseline firing rates (p>0.05) (Fig 2B). More OFC units in CIE mice (67%) than Air mice (54%) showed significantly altered firing rates during any decision-making epochs (Figure 2C) (χ^2^_1,667_ = 10.21, p < 0.002) (see Materials and Methods). However, in both groups, we found similar proportions of OFC units that significantly up-modulated (increased firing rate) or down-modulated (decreased firing rate) across different decision-making epochs (χ^2^’s < 3.76, ps’ > 0.05) (Figure 2D). Furthermore, we found that in both Air and CIE mice, individual OFC units usually altered activity across multiple epochs (e.g. both the onset and offset of a lever press) (Figure 2E) (χ^2^_6,394_ = 8.04, p < 0.24), with relative high percentages of OFC units encoding action onset, action offset, and reward (Air = 18%; CIE = 23%). Altogether, this suggests that the OFC populations normally recruited during this instrumental task were largely not altered following the induction of alcohol dependence.

### Prior CIE procedures enhances OFC action-related activity

While we observed similar recruitment of OFC populations during behavior, a more likely effect of CIE procedures given prior working showing increasing OFC activity is sufficient to restore decision-making (Renteria et al., 2018), would be on the magnitude and patterns of OFC activity during decision-making. As prior work has shown OFC activity modulation related to lever pressing during decision-making in mice (Gremel & Costa, 2013), we first asked whether CIE would alter OFC activity associated with the initiation of actions. We first examined the averaged z-scored activity changes from baseline of significantly modulated units during the 1000-ms period before the lever press onset. The observed changes in firing rates prior to lever pressing were reflected in the normalized activity peri-event heatmaps in both Air and CIE animals (Figure 3A). We found greater increases in OFC firing rates prior to the onset of lever pressing in CIE mice compared to Air mice (Figure 3C). A two-way repeated measures ANOVA (250-ms Bin x Treatment) of significantly modulated unit activity prior to lever press onset showed no interaction, but a main effect of Treatment only (F(1, 1032) = 17.39, p<0.0001). This increase in firing rate was present in both up- and down-modulated CIE population analyses (Figure 3 — figure supplement 1A and 1B), suggesting an overall increase in action-related OFC activity in CIE mice.

**Figure 3.**
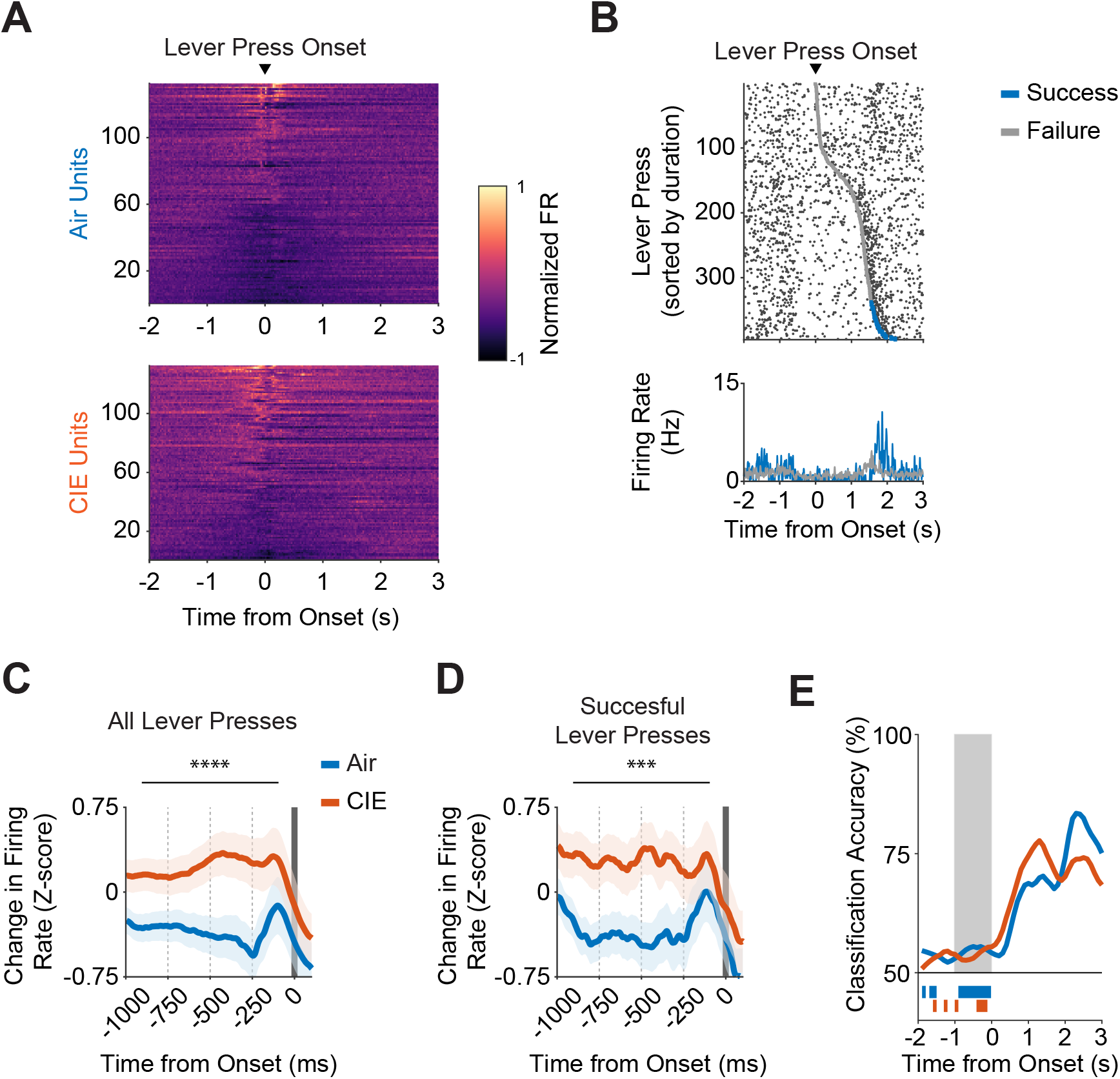
CIE alters OFC activity correlates of action initiation. **(A).** Heat map of normalized firing rates for units that significantly increased or decreased from baseline, displayed relative to lever press onset. Units are sorted by activity from a 1 second window around lever press onset. **(B).** Raster plot of a representative unit’s firing rate relative to lever press onset, sorted from shortest to longest durations within a 1600-ms lever press criterion session that occurred late in training. Grey and blue markers indicate the end of lever presses that failed or succeeded in exceeding the 1600-ms lever press criterion, respectively. **(C).** Average z-scored firing rate changes from baseline for all lever presses and **(D)** successful lever presses only. Firing rate changes were compared across four 250-ms bins relative to lever press onset. **(E).** SVM classification accuracy of task performance outcomes (i.e. lever press successfully exceeded 1600-ms hold-down criterion) from all captured Air and CIE units, displayed relative to lever press onset. Bars underneath traces indicate time points before the onset of the lever press in which classification accuracy was significantly different compared to the null distribution. Shaded region indicates time points in which classification accuracy comparisons were made between Air and CIE groups. Data points represent mean ± SEM. ***p<0.001, ****p<0.0001.

We next asked whether activity associated with lever press initiation reflected future performance outcomes, i.e. were firing rates different for lever presses that were eventually rewarded? To this end, we segmented lever press durations by whether they successfully exceeded the lever press duration criterion or not. For example, the representative unit shown in Figure 3B significantly modulated its firing rate before the onset of a lever press similarly between rewarded and unrewarded lever presses. Indeed, when examining the averaged z-scored population activity changes of significantly modulated units in either Air or CIE mice, we saw no evidence for predictive coding of performance outcomes (Figure 3 — figure supplement 1C and 1D). Given this, the increased firing rate observed in CIE mice was still present when we examined only successful lever presses (two-way repeated measures ANOVA (Bin x Treatment): main effect of Treatment only (F(1, 1032) = 14.58, p=0.0001) (Figure 3D). Furthermore, a support vector machine model trained with the peri-event activity of all captured units to test whether firing rates could accurately classify whether an individual lever press exceeded the 1600-ms duration criterion showed that Air and CIE mice showed similar low classification performance. A two-way repeated measures ANOVA (Bin x Treatment) comparisons of temporally binned classification performance between Air and CIE mice revealed no significant differences within the 1000-ms period prior to lever press onset (p>0.29). Thus, prior CIE procedures increased OFC activity associated with action onset; however, this activity as well as activity in Air control mice was not predictive of impending lever-press success.

### Prior CIE procedures have little effect on OFC activity during action execution

Recent hypotheses have suggested a role for OFC activity during inference of unobservable information needed to support decision-making (Stalnaker et al., 2014; Zhou et al., 2019). We next asked whether the increased activity observed in CIE mice before lever-press onset would persist as they held down the lever. This lever press task used duration as the action contingency, and mice produced a distribution of durations. However, it remained unclear what patterns of OFC activity to expect during the prolonged execution of a lever press, or whether OFC activity would be at similar levels across all durations.

To answer whether increased activity persisted in CIE mice during the duration of the lever press, our analysis focused solely on units that were significantly modulated before the onset of a lever press. When we examined individual unit activity raster plots, we observed broad reductions in OFC activity as mice held down the lever, relative to lever press onset (Figure 4A). Indeed, only an average of ~38% of total lever presses across these significantly modulated OFC units had at least one action potential occur during execution, and this was not different in CIE mice (unpaired t test with Welch’s correction: t_257.4_ = 0.15, p=0.88) (Figure 4B). When these action potentials occurred, they were differentially distributed throughout the lever press. To assess whether CIE procedures altered this distribution, we divided each lever press into four equal duration segments and calculated the average proportion of spikes that occurred within each individual segment (see Materials and Methods). While Air mice showed a U-shaped distribution of activity across the duration of a lever press, this was only slightly altered in CIE mice (Figure 4C) (two-way repeated measures ANOVA (Segment x Treatment); interaction: F(3, 1032) = 2.985, p=0.03; main effect of Segment: F(3, 1032) = 28.08, p<0.0001). In Air mice, the first three segments differed from the last (post-hoc corrected ps’ < 0.0001), while in CIE mice only the 2^nd^ and 3^rd^ segment differed from the last (post-hoc corrected ps’ < 0.0001). In addition, we found that OFC units from CIE mice had a smaller proportion of their spikes occur within the last 25% of their lever press duration compared to their Air counterparts (p=0.05 post-hoc corrected). Thus, OFC did not maintain activity changes during the execution of the duration contingency, and this did not greatly differ between alcohol dependent and control mice.

**Figure 4.**
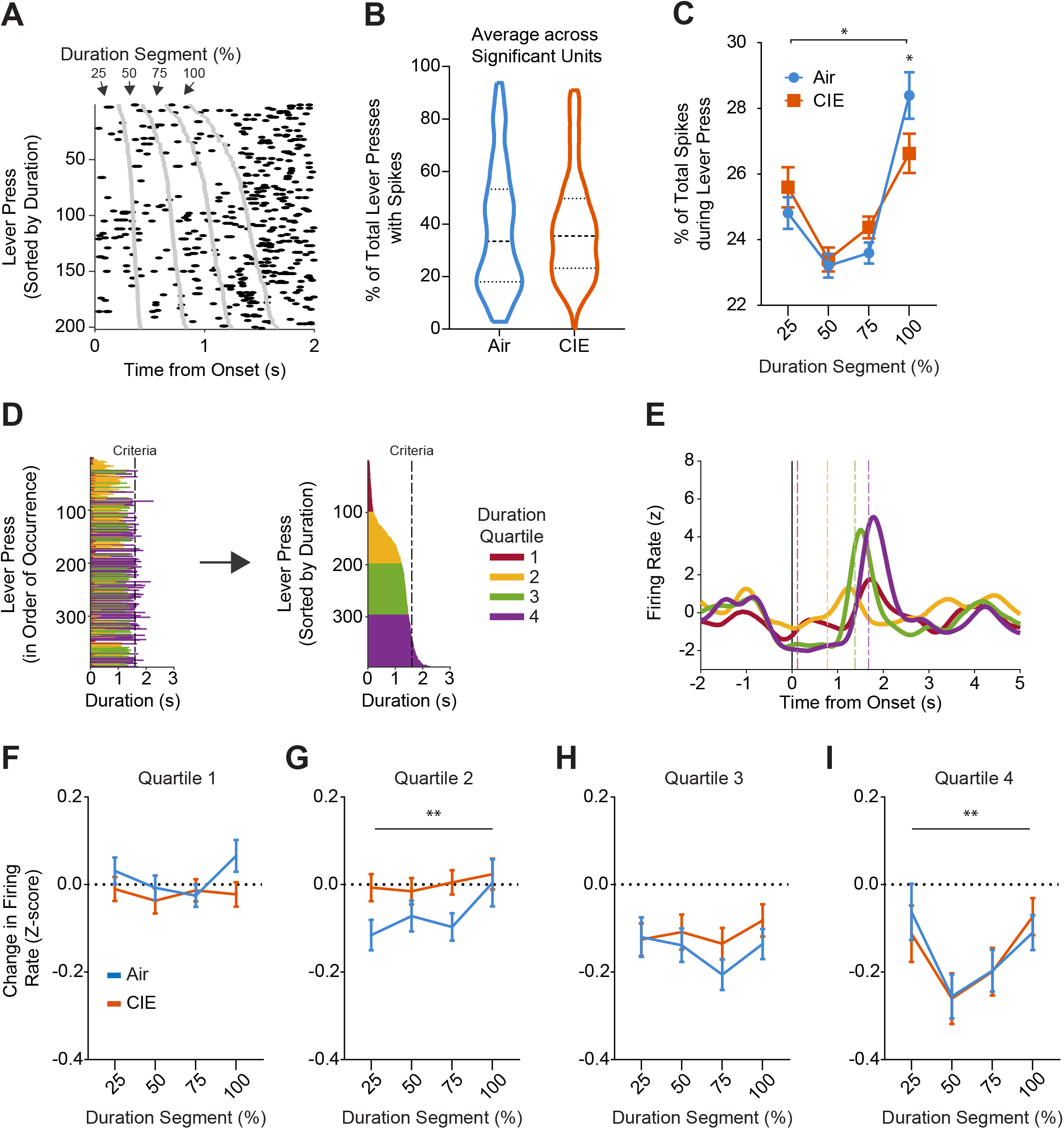
CIE does not alter OFC activity dynamics during actions. **(A).** Raster plot of a representative unit’s firing rate during lever pressing, displayed relative to lever press onset and sorted from shortest to longest durations within a late 1600-ms lever press criterion session. Gray markers indicate the boundaries of equidistant lever press duration segments (i.e. 25%, 50%, 75%, 100% of lever press duration). **(B).** Average percentage of total lever presses within a session during which at least one action potential was detected. **(C).** Average percentage of total spikes that occurred during a lever press, segmented by four equidistant lever press duration segments (i.e. 0-25%, 25-50%, 50-75%, 75-100% of lever press duration). **(D).** Lever presses were segmented into four quartiles determined by the distribution of lever press durations within each individual session. **(E).** Representative unit’s mean z-scored firing rate changes displayed relative to lever press onset. Dashed lines indicate the average lever press duration for each respective duration quartile within that session. **(F).** Mean z-scored firing rate changes from baseline across equidistant duration segments of lever presses belonging to the 1st, **(G)** 2nd, **(H)** 3rd, and **(I)** 4th duration quartiles. Data points represent mean ± SEM. * p<0.05, **p<0.01.

However, the magnitude of OFC activity reductions during ongoing lever presses may depend on the future, total duration of the lever press, and CIE could alter the magnitude of these activity patterns. To examine this, we first segmented each lever press duration and its associated spike activity into four equal duration segments, and then grouped them into duration quartiles determined by the within-session lever press distributions (see Materials and Methods). Then, within each quartile, we examined z-scored activity during the lever press, separated into four equal segments (e.g., 0-25%, 25-50%, 50-75%, 75-100%). Note that these segments are equal in relative terms, though not in absolute duration (Figure 4D and 4E). Quartile distribution boundaries were similar between groups (Figure 4 — figure supplement 1A-1C). This allowed us to make treatment group comparisons of OFC spiking activity during lever pressing, with regards to the future, total duration of that lever press.

We found OFC firing rates during the execution of the lever press did differ depending on the duration of the lever press; however, prior CIE exposure had very little effect on these patterns. As exemplified by Figure 4E, longer lever presses appeared to show lower firing rates during the lever press, with an increase in firing rate occurring close to the release of the lever press. Lever presses in the fourth quartile of duration distributions showed a firing rate that differed as mice held down the lever (two-way repeated measures ANOVA (Segment x Treatment); no interaction; main effect of Segment: F(3, 1032) = 4.939, p=0.002). There was a small, yet significant Treatment difference in the second quartile (two-way repeated measures ANOVA (Segment x Treatment); no interaction; main effect of Treatment: F(1, 1032) = 7.788, p=0.005), but no other significant effects (i.e., no modulation across segments). It is important to note that not all lever press durations in the fourth quartile were rewarded, and that the firing rates measured occurred during the lever-press itself, before the outcome of the lever press was known. Thus, OFC activity during the lever press appears to change in magnitude in relation to the future duration, with longer lever presses showing greater reductions in firing rate. Further, only in the longer lever presses (i.e. those belonging to the 4^th^ duration distribution quartile) was firing rate modulated differentially across the duration of the press. Indeed, the U-shaped pattern of activity during these long lever presses raises the hypothesis that these patterns may reflect an expectancy or confidence signal of future success.

### Prior CIE procedures decrease OFC activity during outcomes

OFC neurons have long been reported to increase their activity in anticipation of and during outcome delivery (Wallis, 2007; Stalnaker et al., 2014, 2018). In the present task, reward delivery cannot occur until the lever is released. Thus, we defined an action offset epoch (1000-ms), and in some cases an outcome-related epoch (3000-ms) following a reward delivery, that encompassed moving to the food receptacle and potentially reward consumption. As the reward is readily visible without mice having to insert their heads into the food receptacle, it is likely that reward perception happens earlier than consumption. As seen in the normalized activity peri-event heatmaps (Figure 5A) and in an example from a representative OFC unit (Figure 5B), OFC firing rates changed significantly during the action offset and outcome-related epochs of the task. Indeed, we observed similar percentages of OFC units recorded between Air and CIE mice where significant activity changes were associated with action offset only (Air = 13.5%, CIE = 16.5%) and outcome-related only (Air = 11.5%, CIE = 7.5%), as well as OFC units that had activity associated with both action offset and outcome evaluation (Air 12%, CIE 7%) or action onset, action offset, and outcome-related (Air = 18%; CIE = 23%) (Figure 2E). Given this, we hypothesized that during action offset and outcome-related epochs, we would observe greater increases in OFC activity in CIE as compared to Air animals, similar to what we observed at action onset. However, we found very little difference in the magnitude of OFC activity or its patterns during the ongoing execution of lever presses between groups. Thus, alcohol dependence may have differential effects on action and outcome-related OFC activity.

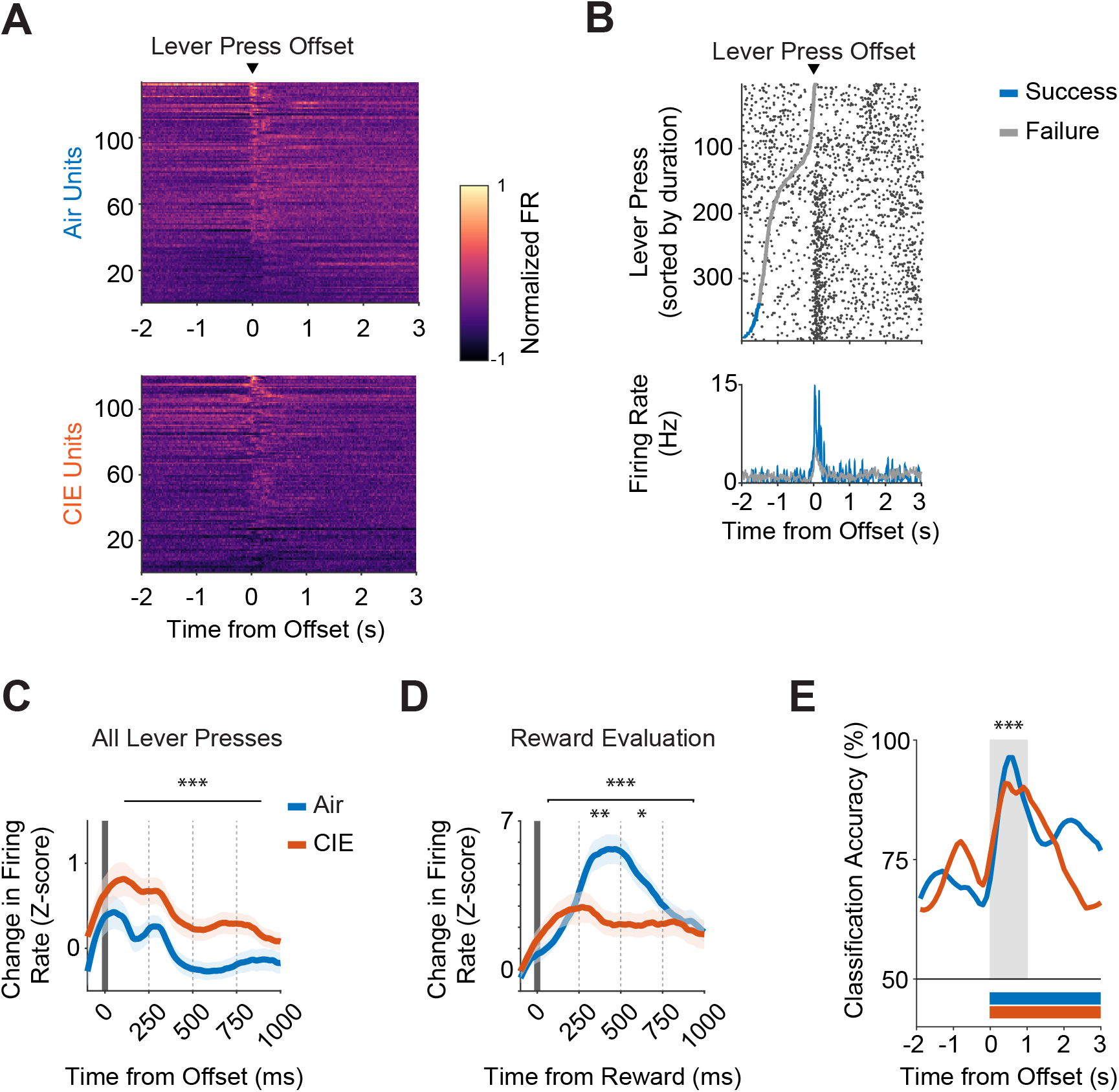
**(A).** Heat map of normalized firing rates for units that significantly increased or decreased from baseline, displayed relative to lever press offset. Units are sorted by activity from a 1 second window around lever press offset. **(B).** Raster plot of a representative unit’s firing rate relative to lever press offset, sorted from shortest to longest durations within a 1600-ms lever press criterion session that occurred late in training. Grey and blue markers indicate the start of a lever presses that failed or succeeded in exceeding the 1600-ms lever press criterion, respectively. **(C).** Average z-scored firing rate changes from baseline for all lever presses and **(D)** rewarded lever presses only. Firing rate changes were compared across four 250-ms bins relative to lever press offset. **(E).** SVM classification accuracy of task performance outcomes (i.e. lever press successfully exceeded 1600-ms hold-down criterion) from all captured Air and CIE units, displayed relative to lever press offset. Bars underneath traces indicate time points after the offset of the lever press in which classification accuracy was significantly different compared to the null distribution. Shaded region indicates time points in which classification accuracy comparisons were made between Air and CIE groups. Data points represent mean ± SEM. * p<0.05, **p<0.01, ***p<0.001

When we examined OFC activity following action offset, we found greater increases in z-scored OFC firing rates in CIE mice compared to Air controls (two-way repeated measures ANOVA (Bin x Treatment); no interaction; main effect of Bin: F(3, 1044) = 6.612, p=0.0002; main effect of Treatment: F(1, 1044) = 22.63, p<0.0001) (Figure 5C and Figure 5 — figure supplement 1A and 1B). We also found that OFC activity changes at action offset reflected performance outcomes in both Air and CIE mice (Figure 5 — figure supplement 1E and 1F); however, it is important to note that this OFC activity was comprised of all lever presses, including those that were rewarded. Thus, we examined OFC firing rate changes during outcome-related epochs following successful lever presses and asked whether CIE procedures would change the magnitude of these increases. We found large increases in OFC firing rate changes during outcome-related epochs. Surprisingly, we found OFC units in Air mice had greater increases in firing rate during outcome-related epochs compared to CIE mice (two-way repeated measures ANOVA (Bin x Treatment); interaction: F(3, 724) = 4.04, p=0.007; main effect of Bin: F(3, 724) = 4.43, p=0.004; main effect of Treatment: F(1, 724) = 8.66, p=0.003) (Figure 5D and Figure 5 — figure supplement 1C and 1D). Given the robust increases in OFC activity during outcome-related epochs, as well as the differences in magnitude from CIE procedures, we asked whether a support vector machine model trained with the peri-event activity of all captured OFC units could accurately classify whether an individual lever press exceeded the 1600-ms duration criterion during action offset and outcome-related epochs. We found high classification accuracy during the outcome-related epochs, especially within the first 1000-ms of reward delivery. Furthermore, classification accuracy was lower in OFC units from CIE mice during this period (two-way repeated measures ANOVA (Bin x Treatment); interaction: (F(9, 180) = 3.33, p=0.0009; main effect of Bin: F(9, 180) = 5.11, p<0.0001). Thus, CIE mice show greater OFC activity associated with lever pressing, but reduced OFC activity during outcome-related epochs, with that activity being less predictive of rewarded decision-making.

## Discussion

Alcohol dependence is associated with impairments to OFC function and aberrant decision-making, thereby increasing the vulnerability to relapse and maladaptive alcohol consumption (Zinn et al., 2004; Chanraud et al., 2007; Loeber et al., 2009; Berre et al., 2012; Reiter et al., 2016; Le Berre et al., 2017). Here, we uncovered neural correlates of actions and outcomes and found them perturbed by prior chronic alcohol exposure and withdrawal. Our results suggest alcohol dependence induces long-lasting perturbations to OFC activity in a bidirectional manner, dependent on the computation being performed. CIE mice showed a modulation of OFC activity that suggests overall increases in OFC activity associated with actions (i.e. lever press onset and offset), and a blunting of OFC activity during outcome-related epochs. This suggests that alcohol dependence does not result in a loss of OFC recruitment, but rather induces a change in how computations performed by OFC circuits may contribute to decision-making.

In the present data, CIE exposed animals acquired and performed the action contingency at similar levels compared to alcohol naïve controls when the value properties of the reward remained stable across training, suggesting that the acquisition of contingency information was unperturbed following alcohol dependence (Figure 1). However, while Air mice adjusted responding following outcome devaluation, CIE mice did so to a lesser degree. Hence our action differentiation task was shown to effectively isolate instrumental learning contingency processes from those crucial for updating and inferring value representations, with the unique feature of doing so in a completely self-paced manner that requires an animal to continuously infer contingency durations to guide their behavior. In contrast to recent findings (Barker et al., 2020), the comparable levels of performance in our task, and the differences observed following outcome devaluation, suggests that the effects of alcohol dependence on behavioral control might be somewhat limited to those requiring outcome evaluation processes. However, Barker et al. examined degradation of the contingency in contrast to acquisition and performance effects examined here. How and whether action contingency and outcome value processes influence each other were not examined. Furthermore, due to difficulties in female mice maintaining and carrying electrodes and associated head-caps, we were not powered in our in vivo recording experiments to examine whether there were any sex differences in our neural data that could differentially mediate contingency and expected outcome value control (Barker et al., 2010). Our data does suggest however that further exploration into alcohol dependence effects on action contingency control and control by expected outcome value is warranted.

Clinical studies have previously shown that alcohol dependence alters representation of decision-making within OFC circuits, albeit not always in the same manner. The OFC is widely found to be hypoactive in alcohol dependence (Volkow et al., 1994, 1997; Boettiger et al., 2007; Sjoerds et al., 2013; Reiter et al., 2016), but there have also been reports of hyperactivity (Wrase et al., 2002; Tapert et al., 2003; Myrick et al., 2004; Hermann et al., 2006; Myrick et al., 2008; Ernst et al., 2014; Reinhard et al., 2015). We find that to be the case in our data as well, such that alcohol dependence altered decision-making representations in the OFC in a variety of ways. Prior CIE exposure and withdrawal changed OFC computations in a way that suggests an overall increase in activity (Figure 3). This dependence-induced change could suggest an increased contribution of OFC processes to action-related processes. We should emphasize that the action contingency in the present task is the duration of the lever press, and that it is inferred from prior experience. This suggests that alcohol dependence increases OFC activity related to the retrieval and execution of inferred action contingencies. In this context, it is important to note that a hyperactive OFC has also been observed in those with obsessive compulsive disorder and increased activity of OFC neurons during actions has been linked to compulsive action phenotypes (Milad & Rauch, 2012; Pauls et al., 2014; Robbins et al., 2019; Lüscher et al., 2020). Whether this increased OFC activity related to actions that we find in CIE mice plays a role in compulsive phenotypes in alcohol dependence is not currently known.

A hallmark of OFC function is its contribution to reward evaluation and updating, with increases in OFC activity observed during outcome anticipation and presentation (Rolls et al., 1996; Jones et al., 2012; Wallis, 2012; Stalnaker et al., 2014, 2015, 2018). Recent works have shown that inhibition of OFC activity during periods of outcome presentation prevent outcome evaluation and updating (Baltz et al., 2018; Malvaez et al., 2019). Further, recent work in humans has suggested that OFC encodes reward identity expectations (Howard & Kahnt, 2018), which contribute to the generation of prediction errors even when there is no change in value (Stalnaker et al., 2018). Here, we observed large increases in OFC activity during outcome-evaluation periods and this increase was reduced following alcohol dependence (Figure 5). SVM modeling showed reduced accuracy in classifying rewarded lever presses following the induction of dependence. Furthermore, CIE affected OFC computations made only after successful lever presses. Thus, in addition to action-associated OFC activity, the data above strongly suggests that OFC’s contribution to outcome retrieval, evaluation, or identification is altered following alcohol dependence. However, in the current experiments we did not test whether the observed increases and decreases in activity correspond with increased and decreased functional contributions of OFC to decision-making. While OFC often reflects decision-making computations, functional necessity of OFC is often limited to periods in which such information is unobservable (Stalnaker et al., 2015). Our data does raise the hypothesis that when OFC is functionally recruited, alcohol dependence may lead to reduced contribution of OFC during outcome-related behaviors. Nonetheless, the observed bidirectional dependence-induced effects on OFC computations support the hypothesized complexity of alcohol dependence effects on OFC decision-making circuitry. For instance, previous accounts on the modulatory influence of alcohol dependence on OFC activity have differed (Volkow et al., 1994, 1997; Boettiger et al., 2007; Sjoerds et al., 2013; Reiter et al., 2016), which in conjunction with our findings suggests a divergent effect of alcohol dependence that may be dependent on decision-making demands and information in OFC. While the critical role for the OFC in regulating the ability to adjust behavior when outcome value or identity changes has been largely established, here we present new evidence on the specificity of dependence-induced effects on the computations supporting these processes.

The self-paced nature of our task allowed us to investigate the dynamics of OFC computations made during ongoing decision-making that relies solely on internal representations or retrieval of learned duration contingencies, rather than a reliance on predictive external sensory information. Here, we show with in vivo electrophysiology data that the OFC is actively engaged during action differentiation. OFC activity changed throughout individual lever press durations, with activity during long lever presses resembling a U-shaped pattern that suggest a working process for inferring the proximity to achieving a desired goal (Figure 4). Moreover, these continuous patterns of activity at first reflect an overall decrease relative to baseline, that subsequently increases prior to the release of the lever press and prior to when outcomes are expected. The increase in activity prior to lever press release may correspond to a greater confidence in outcome delivery, something previously shown for OFC activity in cued tasks (Kepecs et al., 2008; Masset et al., 2020). Prior work in humans has also suggested that OFC lesions have altered subjective reports of time perception (Berlin et al., 2004). Interestingly, prior alcohol dependence had very minor effects on these activity patterns. That both groups showed OFC modulation prior to (with CIE mice exhibiting a larger increase) and during a lever press suggests that initial information used to guide decision-making is still represented and may be more effective at recruiting OFC activity following alcohol dependence.

The dichotomy of CIE effects on the different behavioral components of our task suggests a combination of OFC circuitry changes that could manifest in a variety of ways. In the future, it will be of interest to examine how CIE perturbs OFC function and output that relies on information received by interconnected structures. For example, chronic alcohol exposure and withdrawal may be perturbing the excitability and transmission of local OFC circuitry via cell-type specific changes (Badanich et al., 2013; McGuier et al., 2015; Nimitvilai, Uys, et al., 2017; Renteria et al., 2018), such that the integration of incoming information from other associative regions necessary to guide decision-making is disrupted. Additional difficulties in parsing the effects of alcohol dependence on decision-making processes arise from a fundamental lack of structural and functional input-output mapping of the highly complex neural circuits that support decision-making. While in general the areas projecting to OFC have been identified, the relative proportions of inputs across brain regions, as well as the connectivity, strength, and pattern of inputs onto excitatory and inhibitory OFC populations is unknown in naïve circumstances, much less following alcohol dependence. It may be that alcohol dependence results in a redistribution of inputs across OFC excitatory and inhibitory populations and/or alters input transmission onto OFC circuits, thereby altering their ability to contribute to decision-making.

As with all brain areas, the capacity to contribute to decision-making computations is going to depend on the afferent inputs as well as local capabilities. Alcohol dependence is likely to affect both across the brain. Here we identified some of the complexity in how OFC’s contributions to decision-making computations are altered following alcohol dependence. These findings will hopefully shed light on the behavioral and OFC-based perturbations previously reported and provide insight into the therapeutic treatment of alcohol dependence.

## Materials and Methods

### Animals

Male and female C57BL/6J mice (n = 15, 9 males, 6 females for non-recording experiments; n = 18, 17 males, 1 female for recording experiments) were housed 2–5 per cage under a 14/10 hour light/dark cycle with access to food (Labdiet 5015) and water ad libitum unless stated otherwise. C57BL/6J (The Jackson Laboratory, Bar Harbor, ME, USA) mice were at least 6 weeks of age prior to intracranial micro-array implant and at least 52 days of age prior to vapor procedures or behavioral training. Investigators were not blind to the experimental groups. The Animal Care and Use Committee of the University of California, San Diego approved all experiments and experiments were conducted according to the NIH guidelines.

### Surgical Procedures

Animals under isoflurane anesthesia were implanted with a stereotaxically guided fixed micro-array consisting of four-rows of four platinum-plated tungsten electrodes (35 μm tip, Innovative Neurophysiology, Durham, NC, USA), with electrodes spaced 150 μm apart, and rows 150 μm apart. The dearth of female mice in the recording study was due to problems with female mice not being able to maintain and carry the electrode implant through CIE procedures and behavioral testing. To maximize targeting of the OFC, arrays were centered at the following coordinates from bregma: A, 2.5 mm; M/L, 1.3 mm; V, 2.0 mm. An additional bilateral craniotomy was made over the posterior cerebellum for placement of screws wrapped with the electrical reference wire attached to the micro-array. After testing, mice were euthanized, and brains extracted and fixed in 4% paraformaldehyde. Micro-array placement was qualified by examining tracts in 50–100 μm thick brain slices under a macro fluorescence microscope (Olympus MVX10). A subset of micro-arrays was dyed with a 25 mg/ml Dil (1,1’-Dioctadecyl-3,3,3’,3’-tetramethylindocarbocyanine perchlorate) solution in 200 proof ethanol (Sigma) for placement verification. All surgical and behavioral experiments were performed during the light portion of the cycle.

### Chronic Intermittent Ethanol Exposure and Repeated Withdrawal

One to two weeks after micro-array implant surgeries for recording mice, all mice were exposed to four rounds of ethanol vapor or air (H. C. Becker & Hale, 1993; Howard C. Becker & Lopez, 2004; Marcelo F. Lopez & Becker, 2005; Griffin et al., 2009; Renteria et al., 2018). Each round consisted of 16 hours of vapor exposure followed by an 8-hour withdrawal period, repeated for 4 consecutive days. The CIE procedure is designed to repeatedly induce alcohol withdrawal syndrome after long periods of alcohol exposure, a key criterion in the diagnosis of alcohol dependence (Marcelo F. Lopez & Becker, 2005). Ethanol was volatilized by bubbling air through a flask containing 95% ethanol at a rate of 2–3 L/min. The resulting ethanol vapor was combined with a separate air stream to give a total flow rate of approximately 10 L/min, which was delivered to the mice housed in Plexiglas chambers (Plas Labs Inc). Mice were not pre-treated with a loading dose of ethanol or pyrazole to avoid confounding effects of stress that can bias reliance on habitual control, as well as to avoid the effects of pyrazole on neural activity, including actions at the N-methy-D-aspartate (NMDA) receptor (Pereira et al., 1992; Howard C. Becker & Lopez, 2004; Dias-Ferreira et al., 2009). Blood ethanol concentrations (BEC) were collected at the end of each round from separate, non-experimental mice to avoid previously reported stress effects on decision-making from blood extraction on Air mice (mean ± SEM BEC = 29.53 ± 2.36 mM). BEC assays that experienced technical errors were excluded from this measurement.

### Behavioral Task

We adapted a lever press hold down task previously used to assay the timing of decision-making actions in mice (Yin, 2009; Fan et al., 2012). Mice were trained in standard operant chambers with one lever extended to the left (or right) of a food magazine and a house light on the opposite wall within sound-attenuating boxes (Med-Associates). Two days before training, mice were food restricted and maintained at 85%-90% of their baseline body weight throughout training and testing.

#### Magazine training

On the first day, mice were trained to retrieve pellets from the food magazine (no levers present) on a random time (RT) schedule, with a pellet outcome delivered on average every 120 seconds for 60 min.

#### Continuous reinforcement

The next 3 days the left (or right) lever was present the entire duration of the session. Lever presses were rewarded on a continuous reinforcement (CRF) schedule for up to 15 (CRF day 1), 30 (CRF day 2), or 60 (CRF day 3) pellet deliveries or until 60–90 minutes had passed. For electrode-implanted animals, an additional CRF training day (4 days total) was administered with the implant connected to the amplifier board to habituate the animal to the tethered connection.

#### Lever press hold down training

The action differentiation task required lever press durations to exceed a duration criterion assigned prior to the start of the daily session. This criterion was the minimum duration of time the animal was required to hold the lever in a depressed position to receive a reward. Each session began with the house light turning on and the left (or right) lever being extended for the duration of the session. Lever pressing was self-initiated and self-paced without an imposed trial structure (i.e. the lever was never retracted until the session was complete). Reward delivery occurred at the offset of the lever press only if the hold down timer exceeded the session’s assigned duration criterion. Sessions were completed when 30 outcomes (non-recording animals) or 60 outcomes (recording animals) were earned or after 90 minutes, whichever came first. The lever press duration criterion for the first 5 days was 800 milliseconds, followed by 5 days (4 days in 3 animals due to loss of head-cap implant) of a lever press duration requirement of 1600 milliseconds.

#### Devaluation testing

Following the last day of hold down testing in the behavioral cohort, mice were habituated to a novel cage and 20% sucrose solution for an hour each. Devaluation testing through sensory-specific satiation was conducted across 2 days and consisted of a valued day and a devalued day. For the valued day, the mice were allowed to pre-feed for 1 hour on 20% sucrose solution. For the devalued day, mice could pre-feed for 1 hour on the pellet outcome previously earned in the lever press hold down task. Mice that did not consume enough pellets (< 0.1 grams) or sucrose (< 0.1 ml) during pre-feeding were excluded from subsequent analysis (CIE cohort, n = 1). Each day immediately following pre-feeding, mice were placed into their respective operant chamber for 10 min where the number and duration of lever presses made were recorded, but no outcome was delivered. Investigators were not blind to the experimental groups. Valued and Devalued days were counterbalanced and run across consecutive days.

### Electrophysiological Recordings and Spike Sorting

Spike activity and local field potentials were recorded using an RHD2000 USB interface board system connected to an amplifier board via a serial peripheral interface (SPI) cable (Intan Technologies, Los Angeles, CA, USA). Electrode signals were amplified, digitized at 30 kHz and filtered between 0.1 Hz and 6 kHz for spikes and 0.1 Hz and 600 Hz for local field potentials. Initial sorting occurred prior to each testing session using an online-sorting algorithm (OpenEphys (Siegle et al., 2017)). Behavior events that occurred inside the operant boxes were timestamped in synchronization in OpenEphys with neural activity using TTL pulses collected at a 10 ms resolution from Med Associates SuperPort Output cards. Spike data was re-sorted offline (Offline Sorter, Plexon, Inc.) using a T-Distribution Expectation-Maximization Scan algorithm in 3D feature space (Shoham et al., 2003). This allowed for the identification of neuronal activity units based on waveform, amplitude, and inter spike interval histogram (no spikes during a refractory period of 1.4 ms). After sorting, each isolated cluster of waveforms was then manually inspected, and biologically implausible waveform clusters were removed from further analysis. To ensure high signal-to-noise quality of each waveform cluster, waveforms 2 standard deviations greater than the clustered population mean were excluded from the analyses. Units with < 1,000 spike waveforms captured within an entire recording session or that did not show consistent activity across a recording session were not included in our analyses. Before each recording session, mice were exposed to a brief (10-20 second) bout of low-dose isoflurane anesthesia to connect the implant with the recording cable. To avoid confounding effects of anesthesia on brain activity, mice were then moved into the procedure room and monitored for a minimum of 30 minutes before placing them in the operant chamber and initiating the session.

### Identification of Significantly Modulated Units

To initially examine task-related neural activity, for each previously isolated recorded unit we constructed a peri-event histogram (PETH) around time-stamped lever-press and reward delivery events, such that neural activity was binned into 20-ms bins and averaged across events to analyze amplitude and latency during the recorded behaviors. Per-unit PETHs were then smoothed using a Gaussian-weighted moving average over three bins (60 ms). Using the distribution of the PETH from 10,000 to 2,000 ms before lever press onset as baseline activity, we focused our analysis on a period 2,000 ms before to 10,000 ms after task-related events. A task-related neuron was up-modulated if it had a significant increase in firing rate defined as at least 4 bins (80 ms) with a firing rate larger than a threshold of 95% confidence interval above baseline activity during the period from 2,000 before to 3,000 ms after each task event. A task-related neuron was down-modulated if it had a significant decrease in firing rate if at least 4 consecutive bins (80 ms) had a firing rate smaller than a threshold of 95% confidence interval below baseline activity during the period from 2,000 before to 3,000 ms after each task event (Jin & Costa, 2010). The onset of significantly modulated task-related activity was defined as the first of these four consecutive significant PETH bins. To examine the net effect of CIE on OFC activity as animals performed the task, we combined these up- and down-modulated unit populations for subsequent population analyses.

### Population Analyses

#### Performance-related spike activity

To investigate differences in peri-event spike activity between lever presses that were rewarded or not, spike timestamps occurring 10,000 ms before to 10,000 ms after individual lever press events were split into successes (lever press duration exceeded session’s criterion duration) and failures (lever press duration did not exceed session’s criterion duration). Performance segmented neural activity was then binned into 20-ms bins, averaged across events, and then smoothed using a Gaussian-weighted moving average over three bins (60 ms), resulting in two PETHs per unit (successes or failures). Individual PETHs were then converted to z-scores using the mean and standard deviation of the firing rate during a baseline period occurring 10,000 to 2,000 before lever press onset. Per-unit z-scored PETHs were then averaged by treatment group to construct population response profiles for each group. Population spike activity from the last the two sessions of 1600-ms duration criteria was grouped such that a minimum of one session per animal was included. Population spike activity traces were then smoothed with Matlab’s Savitzky-Golay *smoothdata* method using a 400-sample sliding window for visual display purposes only.

#### Ongoing lever press-related spike activity

To investigate differences in spike activity during ongoing lever-presses, each lever press duration was first calculated by subtracting the lever press onset timestamp from lever press offset timestamp. Each lever press duration was then segmented into four equivalent segment bins (i.e. 0-25%, 25-50%, 50-75%, 75-100% of lever press duration), and all spikes occurring within each of these duration bins were counted and calculated as a proportion of all spikes that occurred during that entire lever press.

To investigate differences in firing rate changes between different lever press durations, lever presses were first grouped into four quartiles determined by the distribution of lever press durations within each individual recording session. Quartile-grouped spike activity occurring 10,000 ms before to 10,000 ms after lever press onset was then binned into 20-ms bins, averaged across lever presses, and then smoothed using a Gaussian-weighted moving average over three bins (60 ms), resulting in four PETHs per unit, one for each duration quartile. To account for variable lever press durations, PETHs were converted to z-scores using the mean and standard deviation of activity occurring before the onset of the lever press proportionate to the average lever press duration within each quartile. Individual lever press activity from these PETHs were then segmented into four equivalent segment bins (i.e. 0-25%, 25-50%, 50-75%, 75-100% of lever press duration). Per-unit, baseline z-scored traces were then averaged across the four duration segments within quartile and treatment groups to construct population response profiles.

#### Neural decoding of task performance from spike activity

For all units from recording sessions in which a minimum of 10 lever presses exceeded the session’s lever press duration criterion, spike timestamps occurring 2,000 ms before to 10,000 ms after individual lever press events were binned into 1-ms bins and labeled by lever press outcome (success or failure to exceed the session’s lever press duration criterion). These peri-event rasters were then segmented by treatment groups and task event (lever press onset or offset) and used to train a model to classify successful lever presses. The classifier, a support vector machine model implemented in Matlab with the NDT toolbox, was trained and tested at 100 ms steps with a bin width of 200 ms (Meyers, 2013). For each of these time points, the classifier used 10 cross-validation splits to segment per-unit firing rates from randomly selected lever press events into training (90%) and testing (10%) sets for 500 resampling runs. Significance at each of these timepoints was tested by first creating 5 null distributions of decoding accuracy with 500 resampling runs each in which the performance labels were shuffled. The accuracy of our decoder was then compared to these null distributions across all time points.

### Statistical Procedures

Statistical significance was defined as an alpha of *p* < 0.05. Statistical analysis was performed using GraphPad Prism 8.3.0 (GraphPad Software) and custom MATLAB R2019a (MathWorks) scripts (https://github.com/gremellab/CIEOFCHOLD). Acquisition data, including lever presses, response rate, and proportion of lever presses that were rewarded were analyzed using two-way repeated measures ANOVA (Session x Treatment) unless otherwise noted. For outcome devaluation testing, two-way repeated measure ANOVAs (Value x Treatment) with pre-planned post-hoc Sidak’s multiple comparison testing were performed to examine whether outcome devaluation reduced lever pressing on the devalued compared to valued day within each group. For peri-event spike activity comparisons, per-unit average z-scored firing rates were binned into four 250-ms bins before the lever press onset, or after lever press offset and after reward delivery, respectively. Within treatment groups, we performed two-way repeated measure ANOVAs (Bin × Outcome) to examine differences in spike activity between lever presses that failed or succeeded to exceed the session’s lever press duration criteria, with post-hoc Sidak’s multiple comparison testing to determine bins in which differences were pronounced. Two-way repeated measure ANOVAs (Bin x Treatment) were performed to examine differences in spike activity between treatment groups, with post-hoc Sidak’s multiple comparison testing to determine bins in which differences were pronounced. Two-way repeated measure ANOVAs (Segment × Treatment) were performed to examine group differences in the proportions of spikes occurring between lever press duration segments. When appropriate, mixed-effect analyses were conducted in lieu of repeated measures ANOVAs (e.g. when data points were missing due to loss of implant). Data are presented as mean ± SEM.

## Acknowledgements

This research was supported by R00AA021780, R01AA026077, Whitehall and Brain and Behavior Foundations to C.M.G., F31AA027439 to D.C.S., and NSF GRFP grant number DGE-1650112 to C.C. We thank undergraduate research assistant Mariela Lopez Valencia for technical assistance and Roy Jungay for animal care assistance.

---

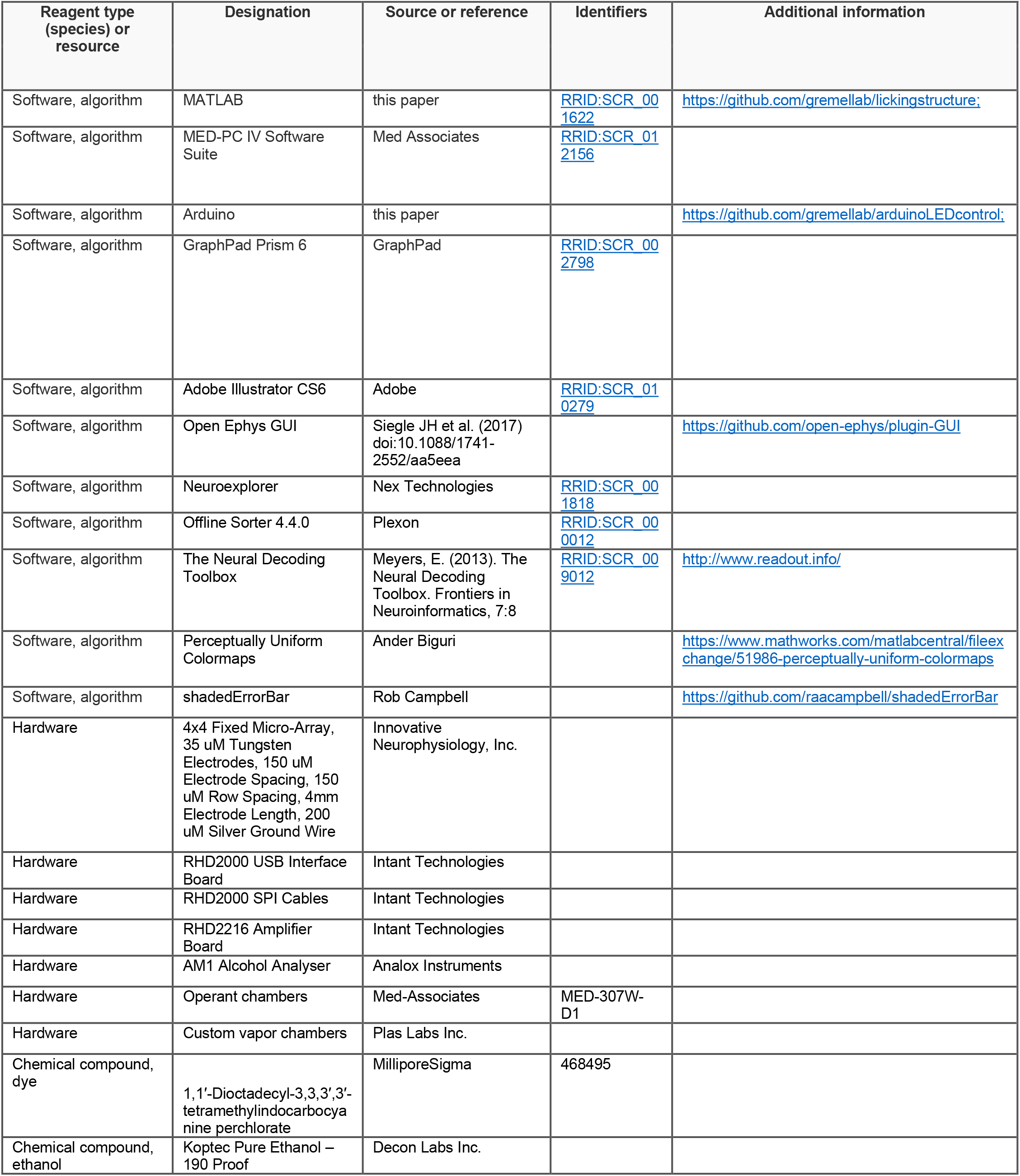
Key Resources Table.

**Figure 1 – figure supplement 1.**
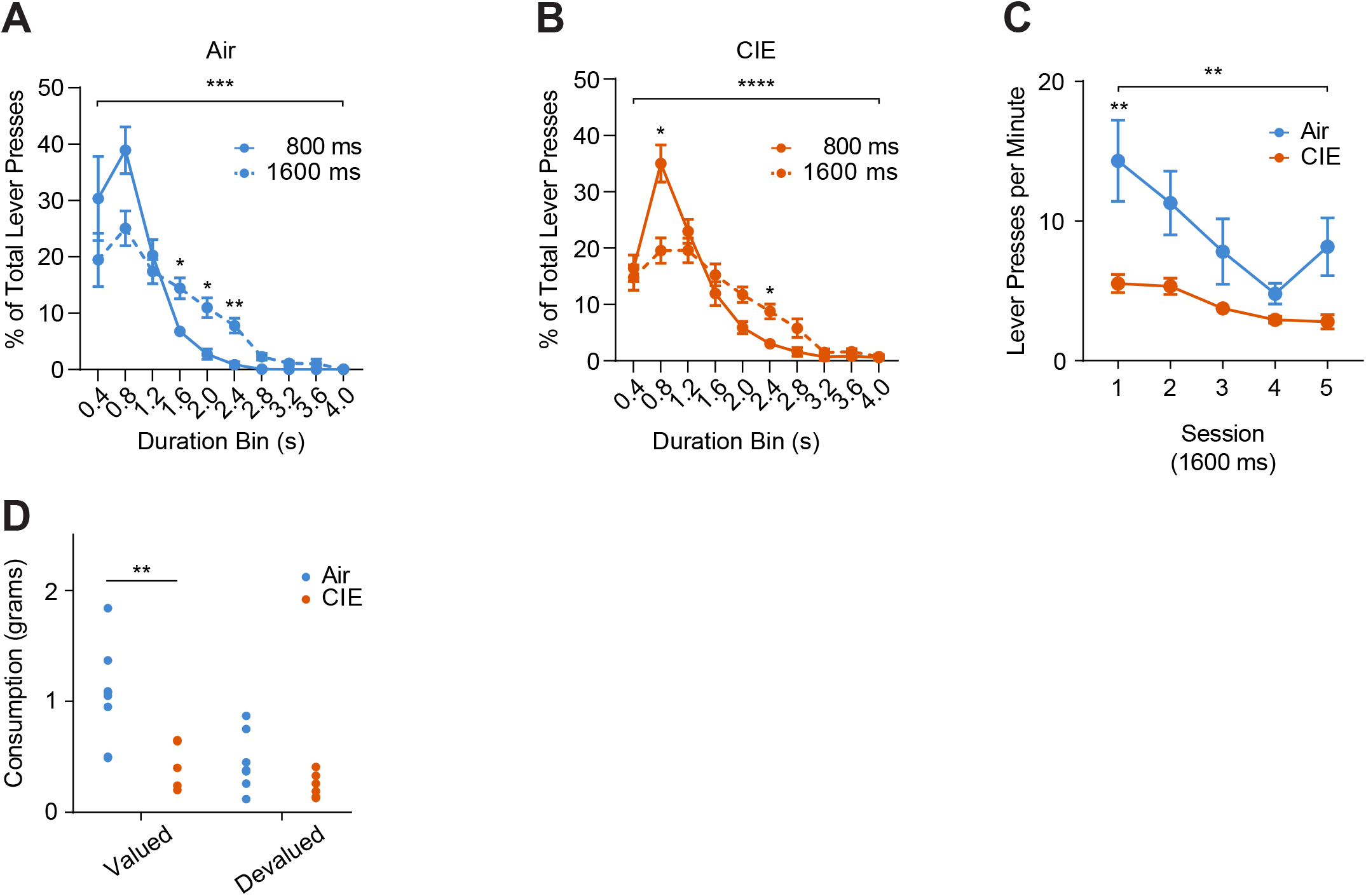
Action differentiation performance throughout acquisition for behavioral cohort. **(A).** Distribution of the percentage of total lever presses binned by duration for Air and **(B)** CIE groups. Distributions were created by averaging across all 5 sessions within each duration criteria. In the Air group, a two-way repeated measures ANOVA (Bin x Criteria) on the lever press duration distribution revealed an interaction: F(9, 126) = 3.91, p=0.0002, and a main effect of Bin: F(9, 126) = 36.89, p<0.0001 and Criteria: F(1, 14) = 4.98, p=0.04. In the CIE group, a two-way repeated measures ANOVA (Bin x Criteria) on the lever press duration distribution revealed an interaction: F(9, 108) = 6.42, p<0.0001, and a main effect of Bin only: F(9, 108) = 56.71, p<0.0001. **(C).** Average lever pressing rate through the 5 daily 1600-ms lever press duration criterion sessions. A two-way repeated measures ANOVA (Session x Treatment) on lever pressing rate found an interaction: F(4, 52) = 4.80, p=0.002; main effects of Session: F(4, 52) = 17.34, p<0.0001 and Treatment: F(1, 13) = 5.83, p=0.03. **(D).** Total grams of food pellet and 20% sucrose solution (Valued state) consumed during the 1-hour ad-libitum access feeding period before each 10-minute 1600-ms duration criterion session. Unpaired t tests with Welch’s correction on total consumption revealed a difference between Air and CIE groups (t10.63=3.398, p=0.0.006) in the Valued state, but not the Devalued state (p>0.09). Data points represent mean ± SEM. *p<0.05, **p< 0.01, ***p<0.001, ****p<0.0001.

**Figure 2 – figure supplement 1.**
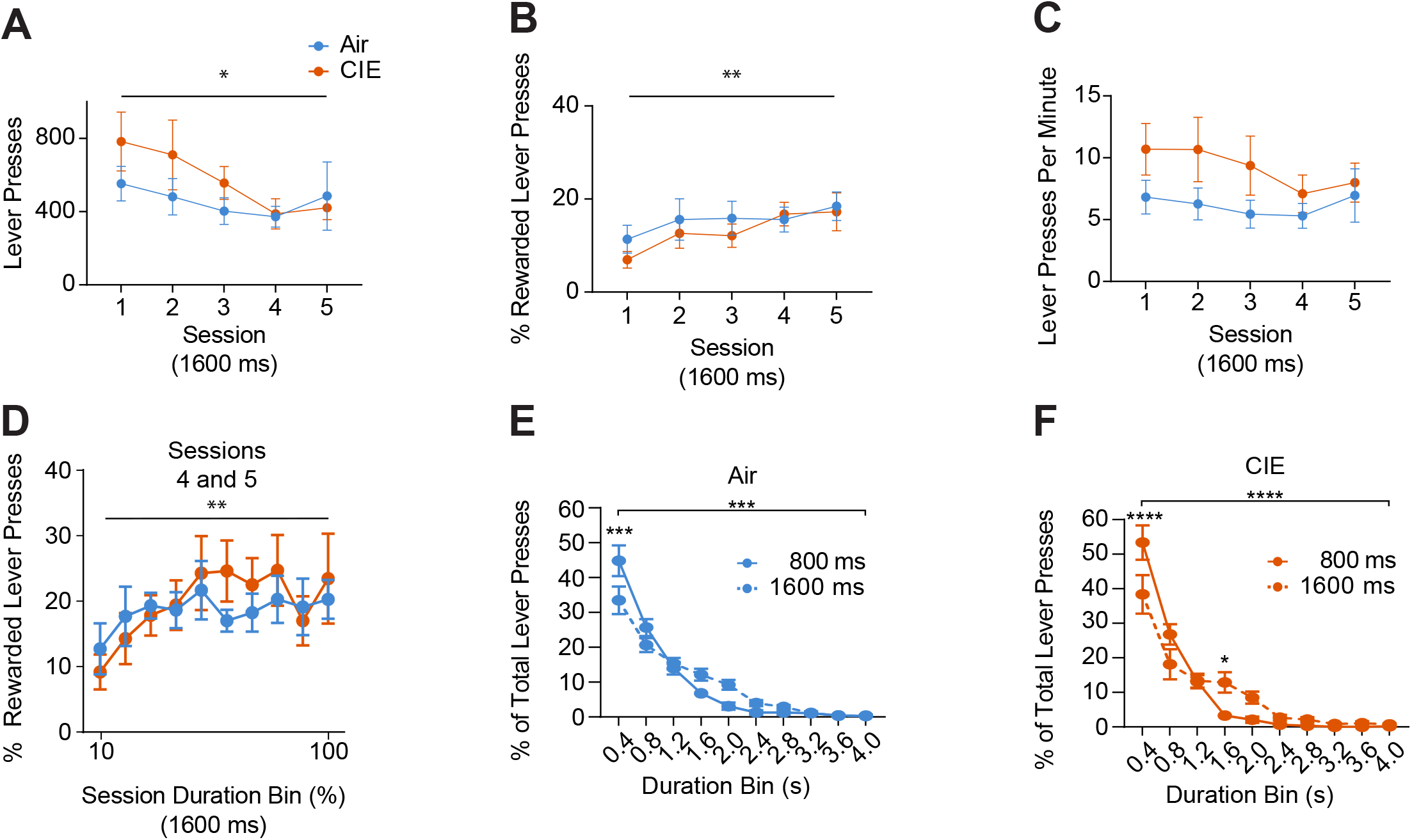
Action differentiation performance throughout acquisition for cohort implanted with chronic indwelling micro-array electrodes. **(A).** Average total lever presses, **(B)** percentage of successful lever presses, and **(C)** lever pressing rate throughout 1600-ms hold down criteria training sessions. A mixed-effects analysis (Session x Treatment) on lever pressing rate throughout 1600-ms duration criterion training sessions found no interaction or main effects (p>0.05). Similar analyses found no interactions, but main effects of Session on the average total lever presses: F(4, 61) = 3.005, p=0.03 and the percentage of successful lever presses: F(4, 61) = 4.369, p=0.004. **(D).** Average percentage of rewarded lever presses within normalized session duration bins during the last two 1600-ms hold down criteria training sessions. A Mixed-effects analysis (Bin x Treatment) found no interactions, but a main effect of Bin only: F(9, 144) = 3.105, p=0.002). **(E).** Distribution of the percentage of total lever presses binned by duration for Air and **(F)** CIE groups. Distributions were created by averaging across all 5 sessions within each duration criteria. A two-way repeated measures ANOVA (Bin x Criteria) on the Air group’s distribution revealed an interaction: F(9, 126) = 3.67, p=0.0004 and a main effect of Bin only: F(9, 126) = 91.28, p<0.0001. A similar analysis on the CIE group’s distribution also revealed an interaction: F(9, 135) = 4.29, p<0.0001, and a main effect of Bin only: F(9, 135) = 73.67, p<0.0001. Data points represent mean ± SEM. *p<0.05, **p< 0.01, ***p<0.001, ****p<0.0001.

**Figure 3 – figure supplement 1.**
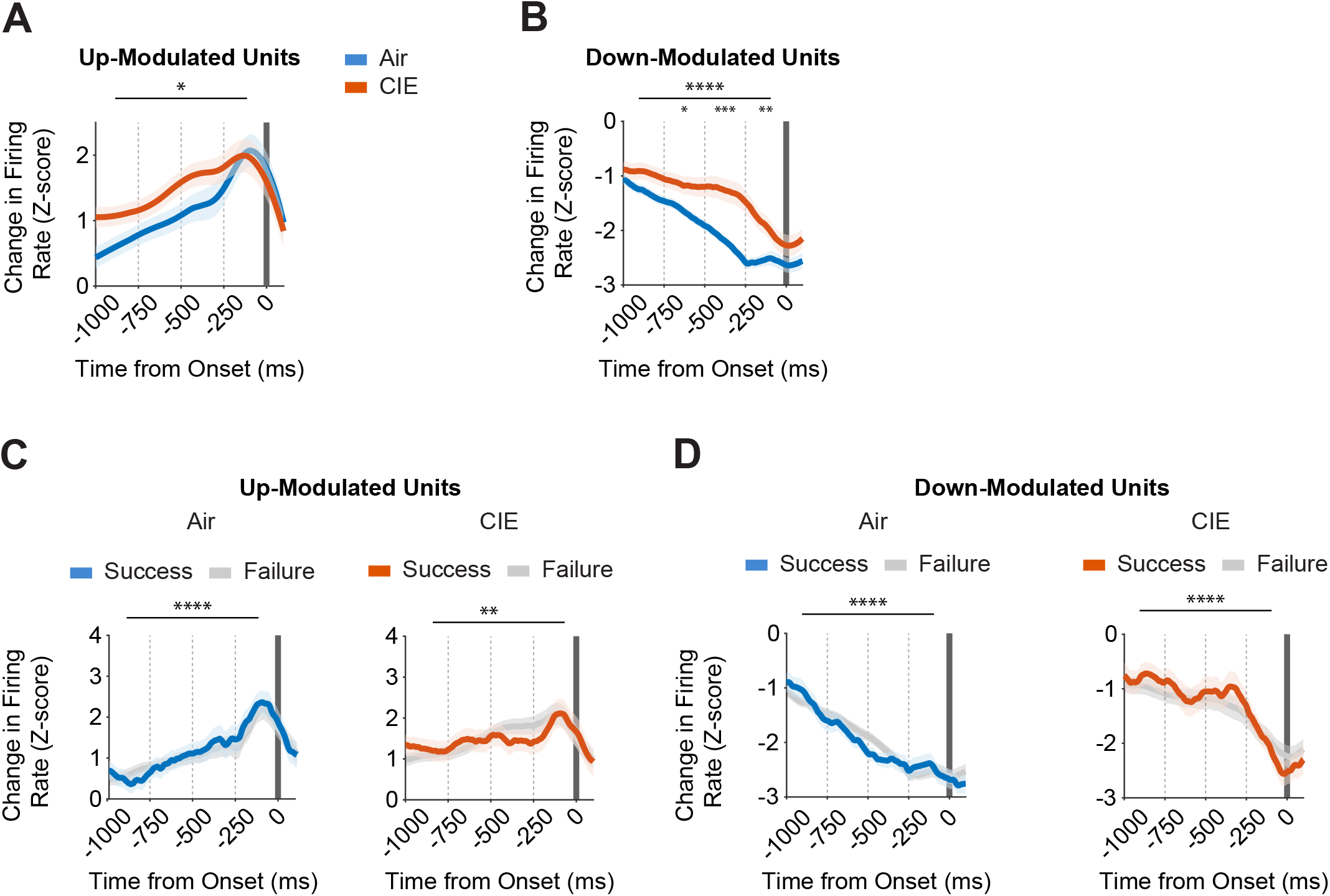
OFC activity correlates of action initiation are altered by CIE but do not reflect future outcomes. **(A).** Average z-scored firing rate changes from all lever presses for significantly up-modulated and **(B)** down-modulated units from Air and CIE groups, displayed relative to lever press onset. In up-modulated units, a two-way repeated measures ANOVA (Bin x Treatment) on activity prior to lever press onset revealed no interactions, but main effects of Bin: F(3, 564) = 12.34, p<0.0001 and Treatment: F(1, 564) = 5.878, p=0.016. In down-modulated units, a two-way repeated measures ANOVA (Bin x Treatment) on activity prior to lever press onset revealed no interactions, but main effects of Bin: F(3, 508) = 25.11, p<0.0001 and Treatment: F(1, 508) = 38.84, p<0.0001. Post-hoc comparisons on down-modulated unit activity found differences between Air and CIE groups in the 2nd (p=0.036), 3rd (p<0.001), and 4th bins (p=0.0045). **(C).** Air (left) and CIE (right) group’s average Z-scored firing rate changes of up-modulated and **(D)** down-modulated units, segmented by whether a lever press successfully exceeded the 1600-ms duration criterion or not, and displayed relative to lever press onset. For the Air group, individual two-way repeated measures ANOVAs (Bin x Outcome) on activity prior to lever press onset revealed no interactions, but main effect of Bin only for up-modulated (F(3, 560) = 14.47, p<0.0001) and down-modulated (F(3, 552) = 37.13, p<0.0001) units. Similarly, for the CIE group, individual two-way repeated measures ANOVAs (Bin x Outcome) on activity prior to lever press onset revealed no interactions, but main effect of Bin only for up-modulated (F(3, 568) = 4.19, p=0.006) and down-modulated (F(3, 464) = 9.77, p<0.0001) units. Average Z-scored firing rate changes were compared between treatment groups or outcomes across four 250-ms bins relative to lever press initiation. Data points represent mean ± SEM. *p<0.05, **p< 0.01, ***p<0.001, ****p<0.0001.

**Figure 4 – figure supplement 1.**
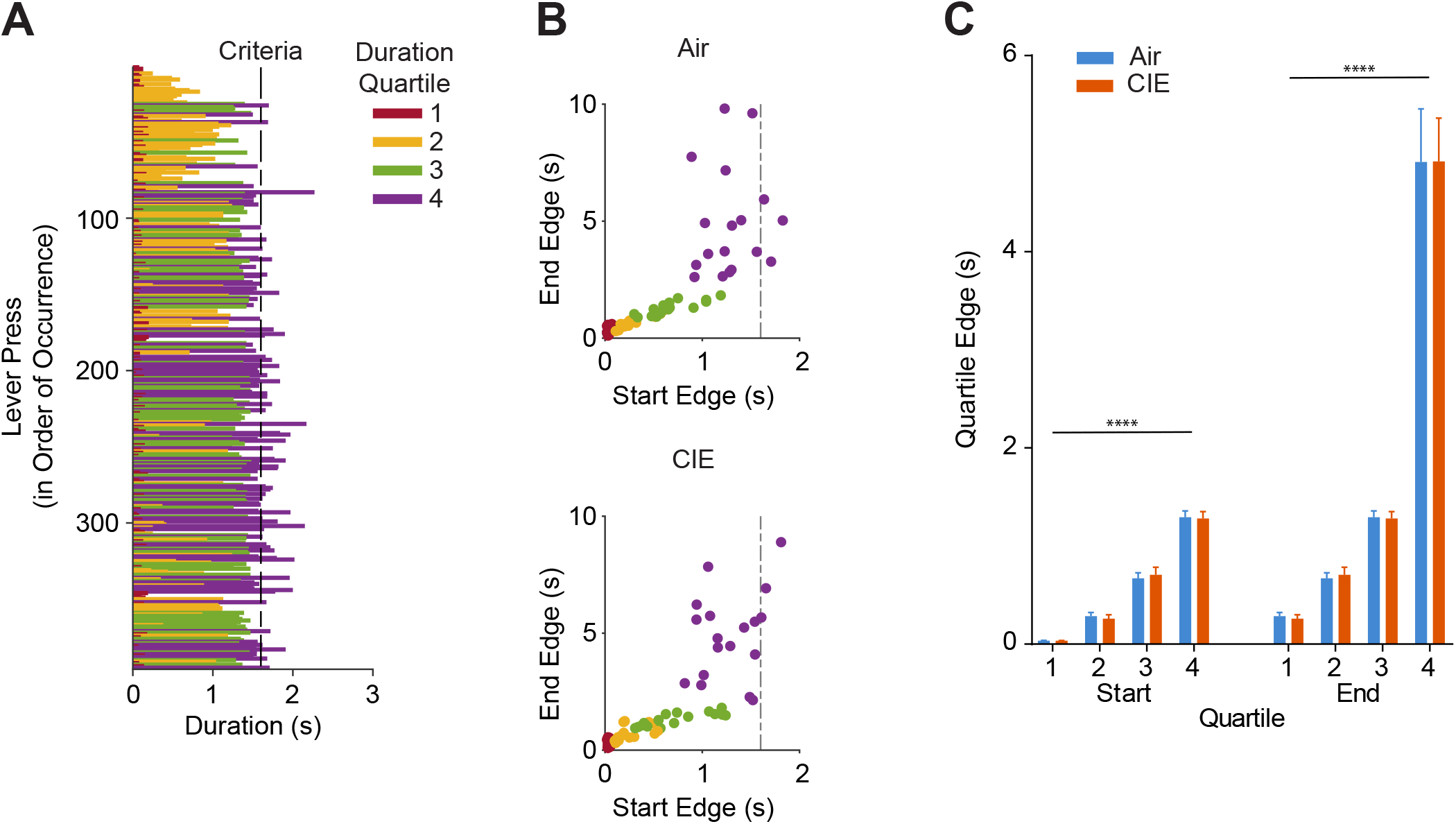
Quartiles were determined by within-session lever press duration distributions and did not differ between Air and CIE groups. **(A).** Examples session in which lever press durations were binned into four quartiles determined by the distribution of lever press durations. **(B-C).** Duration quartile bin boundaries were similar between groups across the last two 1600-ms hold down criteria sessions that were included in our firing rate analyses. For the starting edges of these bin boundaries, a two-way ANOVA (Quartile x Treatment) found no interaction, but a main effect of Quartile only: F(3, 136) = 226.5, p<0.0001). For the ending edges of these bin boundaries, a two-way ANOVA (Quartile x Treatment) found no interaction, but a main effect of Quartile only: F(3, 136) = 141.8, p<0.0001). Data points represent mean ± SEM. ****p<0.0001.

**Figure 5 – figure supplement 1.**
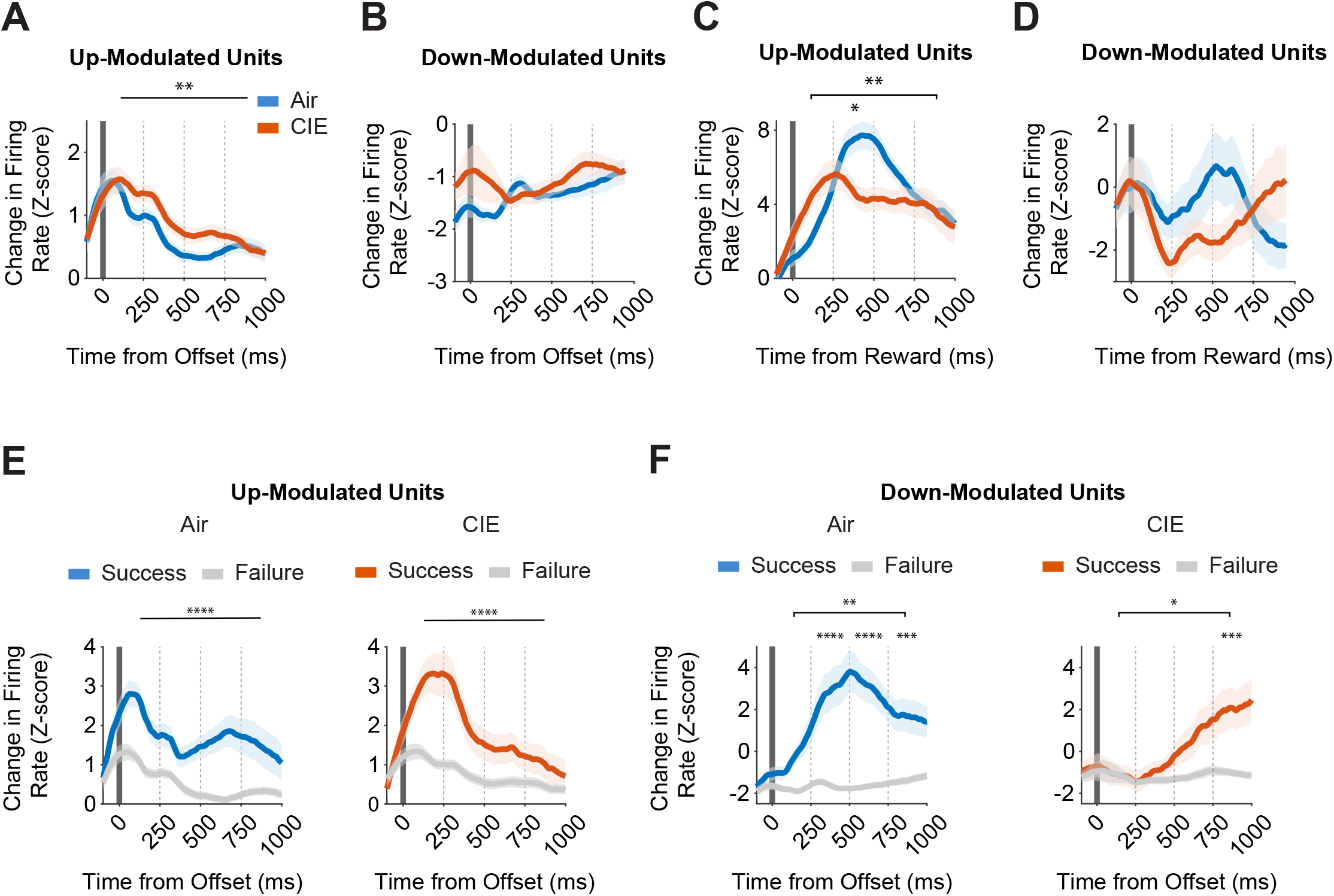
OFC activity correlates during outcome-related epochs are altered by CIE and reflect successful performance. **(A).** Average z-scored firing rate changes from all lever presses for significantly up-modulated and **(B)** down-modulated units from Air and CIE groups, displayed relative to lever press offset. In up-modulated units, a two-way repeated measures ANOVA (Bin x Treatment) on activity prior to lever press offset revealed no interactions, but main effects of Bin: F(3, 748) = 21.82, p<0.0001 and Treatment: F(1, 748) = 8.159, p=0.004. In down-modulated units, a two-way repeated measures ANOVA (Bin x Treatment) on activity prior to lever press offset revealed no differences (p>.08). **(C).** Average z-scored firing rate changes from rewarded lever presses for significantly up-modulated and **(D)** down-modulated units from Air and CIE groups, displayed relative to reward delivery. In up-modulated units, a two-way repeated measures ANOVA (Bin x Treatment) on activity prior to reward delivery revealed an interaction: F(3, 532) = 4.508, p=0.004, and a main effect of Bin: F(3, 532) = 7.199, p<0.0001. Post-hoc comparisons on up-modulated unit activity found differences between Air and CIE groups in the 2nd (p<0.03) bin. In down-modulated units, a two-way repeated measures ANOVA (Bin x Treatment) on activity prior to reward delivery revealed no differences (p>0.2). **(E).** Air (left) and CIE (right) group’s average Z-scored firing rate changes of up-modulated and **(F)** down-modulated units, segmented by whether a lever press successfully exceeded the 1600-ms duration criterion or not, and displayed relative to lever press offset. For up-modulated units, individual two-way repeated measures ANOVAs (Bin x Outcome) on activity after to lever press offset revealed no interactions, but main effects of Bin: F(3, 736) = 5.406, p=0.0011, and Outcome: F(1, 736) = 46.77, p<0.0001 for the Air group and no interactions, but main effects of Bin: F(3, 760) = 8.816, p<0.0001, and Outcome: F(1, 760) = 31.08, p<0.0001 for the CIE group. For down-modulated units, individual two-way repeated measures ANOVAs (Bin x Outcome) on activity after to lever press offset revealed an interaction: F(3, 408) = 3.916, p=0.009, and main effects of Bin: F(3, 408) = 4.523, p=0.004, and Outcome: F(1, 408) = 81.83, p<0.0001 for the Air group, and an interaction: F(3, 248) = 3.412, p=0.02, and main effects of Bin: F(3, 248) = 4.540, p=0.004, and Outcome: F(1, 248) = 12.79, p=0.0004 for the CIE group. Post-hoc comparisons on down-modulated unit activity found differences between successful and failed lever presses in the 2nd (p<0.0001), 3rd (p<0.0001), and 4th (p=0.0002) bins for the Air group and in the 4th (p=0.0002) bin only for the CIE group. Average Z-scored firing rate changes were compared between treatment groups or outcomes across four 250-ms bins relative to the end of the lever press or reward delivery. Data points represent mean ± SEM. *p<0.05, **p< 0.01, ***p<0.001, ****p<0.0001.

